# Translating Music to Touch: Exploring Tactile Perception of Pitch, Roughness, and Pleasantness

**DOI:** 10.1101/2025.09.30.679532

**Authors:** Jeremy Marozeau, Alice Taylor, Sasha Novozhilova, Mario Prsa, Daniel Huber

## Abstract

Music is a rich multisensory experience, yet individuals with hearing impairments often lack access to this important aspect of culture. As tactile technologies advance, there is growing interest in whether musical information can be conveyed through vibration. This study investigates how core dimensions of auditory music perception, pitch, roughness, and pleasantness, can be translated into the tactile domain. Participants were asked to rate these perceptual dimensions in response to sinusoidal and complex waveforms, including amplitude-modulated signals, sawtooth, and missing fundamental stimuli. Perceived pitch showed a systematic relationship with stimulus frequency for most participants, suggesting that tactile devices could at least partially convey some form of simple melodic patterns. The sawtooth waveform emerged as particularly effective for representing pitch changes, underscoring the role of rapid temporal transitions in tactile pitch encoding. Roughness ratings were negatively correlated with pleasantness, mirroring well-established findings in auditory perception. Waveforms with sudden temporal changes or rapid amplitude modulations were consistently judged as less pleasant. Taken together, these findings in normal hearing participants may inform the design of vibrotactile displays that could support access to selected music relevant perceptual dimensions, although generalization to people with hearing loss remains to be tested. Importantly, our results highlight the crucial role of the fast temporal envelope rather than the temporal fine structure, in shaping vibrotactile perception. Consistent with this interpretation, our data provide evidence for a tactile analog of the auditory missing fundamental phenomenon, reinforcing the idea that tactile pitch perception primarily relies on envelope periodicity rather than the presence of specific frequency components.

## I. Introduction

Can music be perceived through skin vibration? For the millions of people with hearing loss who are excluded from the auditory experience of music, haptic technologies offer a potential solution into this essential form of human expression. However, a key question remains: how much of music’s richness can truly survive the translation of sound to touch? These questions have gained interest in recent years with the development of new devices aimed at helping hearing impaired people experience music through tactile stimuli [1]–[5]. Sound and tactile vibrations are inherently linked as both involve the transmission of mechanical waves. It makes the sense of touch a natural medium for conveying musical qualities when auditory processing is impaired [6], [7].

With the rapid improvement of haptic technologies, a wide range of devices has been proposed to convey music through vibrations, see for example the reviews by [8], [9]. Existing solutions rely on different types of actuators, including voice coils, eccentric rotating mass motors (ERM), piezoelectric actuators, and air-based stimulation [10]. Voice coil and piezoelectric actuators are commonly used in experimental and artistic contexts because they offer relatively wide bandwidth and precise control of vibration parameters, while ERM are frequently adopted in wearable and consumer devices due to their low cost and robustness [4]. Each of these technologies presents specific advantages and limitations, and differs in terms of energy efficiency, maximum displacement, acceleration capabilities, and usable frequency range. For instance, a major limitation of commonly used single ERM actuators is that they are controlled by a single parameter, the rotation speed, which inherently couples vibration frequency and intensity, thereby restricting independent control of key perceptual dimensions [8].

In addition to actuator type, haptic music systems vary widely in spatial configuration. Some approaches rely on a single actuator placed on a fingertip or the torso to convey global rhythmic or low frequency information [2], [11], while others distribute multiple actuators around a body part, such as the wrist or embedded in gloves, to exploit spatial patterns of vibration [12]. At the largest scale, full body systems such as vibrating chairs or platforms have been developed to immerse the listener in music through whole body vibration, particularly in concert or installation settings [13], [14]. While each of these approaches offers distinct benefits and constraints, a principled comparison between them requires a deeper understanding of which musical features can be meaningfully transmitted through the skin.

Music is a complex auditory signal that depends on multiple physical parameters such as frequency, intensity, and spectral envelope. These parameters are fundamental in shaping perceptual dimensions, such as pitch, loudness, and timbre, which are low level aspects of music perception. Beyond these basic attributes, higher level perceptual dimensions, including pleasantness, roughness, dissonance, and valence, play critical roles in shaping the emotional and aesthetic experience of music [15]–[17]. For example, pitch height and pitch strength contribute to melody perception, while roughness and dissonance are essential to create tension and resolution, two key elements of musical expression [16], [17]. Translating these auditory features into tactile experiences requires an understanding of how vibration affects the perceptual and emotional aspects of touch.

Auditory pitch is clearly defined by the American National Standards Institute as *that auditory attribute of sound according to which sounds can be ordered on a scale from low to high* [18]; we could extend this to vibrotactile pitch as *that tactile attribute according to which vibrations can be ordered on a scale from low to high*.

Psychophysical studies of vibrotactile pitch perception consistently demonstrate that frequency discrimination through the skin is markedly coarser than in audition. Across stimulation sites, frequency ranges, and psychophysical paradigms, reported Weber fractions for vibrotactile frequency typically fall between approximately 0.2 and 0.3, whereas auditory pure tone discrimination reaches Weber fractions on the order of 0.003 [12], [19]–[22]. This several orders of magnitude difference highlights a fundamental limitation in the spectral resolution of the tactile system relative to the auditory system. Moreover, vibrotactile frequency discrimination is strongly modulated by stimulus intensity, with higher vibration amplitudes yielding improved discrimination performance [19].

A further critical distinction between auditory and vibrotactile pitch perception is that tactile pitch is not determined by frequency alone. Using frequency discrimination paradigms, Prsa and colleagues demonstrated in both humans and mice that systematic changes in vibration amplitude bias perceived vibrotactile pitch, giving rise to families of equal pitch curves in which physically different frequency and amplitude combinations are perceptually matched [23]. These findings indicate that any attempt to quantify perceived pitch differences in the tactile domain must explicitly account for the influence of stimulus intensity, rather than treating frequency as an isolated perceptual dimension.

Importantly, a large frequency discrimination threshold does not necessarily imply a fundamentally different perceptual step size. Although larger frequency differences may be required to detect a change in vibrotactile stimulation, the subjective magnitude of the perceived difference, once detected, may still be comparable to that experienced in audition. Addressing this distinction requires moving beyond discrimination thresholds to directly quantify perceived pitch differences using absolute magnitude estimation, an approach pioneered by Stevens and widely applied in audition and other sensory modalities to characterize perceptual scaling [24]. In the vibrotactile domain, however, applying such an approach necessitates not only varying stimulus frequency but also systematically controlling stimulus intensity in order to neutralize its confounding influence on perceived pitch, as demonstrated by the amplitude-dependent pitch biases described above.

To investigate this, Experiment 1 asked participants to rate the perceived pitch of sinusoidal stimuli using an absolute magnitude scale in both the auditory and vibrotactile modalities. We hypothesized that, once stimulus intensity was appropriately controlled, the relationship between perceived pitch and physical frequency would exhibit a similar slope across modalities, despite large differences in absolute discrimination thresholds.

While sinusoidal stimuli provide a controlled framework for isolating the relationship between frequency and perceived pitch, they capture only a limited aspect of pitch as it operates in music. Musical sounds are typically complex periodic signals with rich spectral structure, and in such signals pitch perception is shaped not only by waveform periodicity, often associated with pitch chroma, but also by the distribution of energy across the frequency spectrum, which contributes to percepts of tone height and timbral brightness. Consequently, two sounds sharing the same fundamental periodicity can differ substantially in perceived pitch-related qualities as a function of their harmonic content [25]. Understanding pitch perception in ecologically valid musical contexts therefore requires moving beyond pure tones to explicitly consider the roles of spectral composition and temporal structure.

A clear distinction must be made between waveform shape, periodicity, and fundamental frequency, *F*_0_. The waveform specifies the temporal shape of the signal and determines the presence and relative amplitude of its harmonic components, while periodicity refers to the repetition rate of the waveform and is quantified as the inverse of *F*_0_. Different waveforms can thus share the same periodicity while exhibiting markedly different spectra. For example, a sinusoid contains a single frequency component, whereas a sawtooth waveform comprises a dense harmonic series with amplitudes decreasing approximately proportionally with the harmonic number. Any periodic signal can further be decomposed into temporal fine structure, capturing rapid cycle-by-cycle oscillations, and a fast temporal envelope, reflecting instantaneous amplitude fluctuations obtained via the Hilbert transform [26]. Whereas fine structure conveys tonal information, the fast envelope plays a dominant role in shaping perceptual attributes such as timbre and speech intelligibility. Experiment 2 was designed to test the hypothesis that, even when periodicity is held constant, differences in waveform shape and spectral composition systematically influence perceived vibrotactile pitch, reflecting contributions beyond fundamental frequency alone, in a similar way to the auditory domain.

Beyond pitch, musical sounds convey affective and qualitative dimensions, such as tension and dissonance, which are commonly associated with perceptual roughness. In audition, roughness is thought to arise from rapid amplitude fluctuations produced by interactions between unresolved harmonic components. Computational models of auditory roughness, most notably that of Zwicker and Fastl [27], emphasize the role of spectral density and partial spacing in higher frequency regions, rather than the fundamental frequency itself, in shaping roughness perception. Sounds with dense harmonic content and substantial high frequency energy are therefore expected to elicit stronger roughness percepts than spectrally sparse sounds.

In the tactile domain, models of roughness perception under single point stationary stimulation are less established. The duplex theory, proposed by Bensmaia & Hollins [28], suggests distinct mechanisms underlying tactile texture perception. Coarse textures, elements greater than 100 *µ*m, are encoded spatially through slowly adapting type I afferents, whereas finer textures, less than 100 *µ*m, are primarily mediated by rapidly adapting afferents, particularly Pacinian corpuscles. Pacinian corpuscles are highly sensitive to high-frequency vibrations, optimally around 250 Hz, suggesting frequency as an important factor for finer textures. Research has proposed two main coding theories for fine texture: an intensity-based code and a temporal, frequency-based code [28]. Specifically, it has been suggested that tactile roughness perception could result from a weighted sum of the stimulus spectrum by the Pacinian corpuscle’s sensitivity profile around 250 Hz, with higher frequencies receiving greater weighting. Additionally, the irregularity and sharp spatial deviations in texture, as measured by the Hurst coefficient [29], have been proposed as critical factors contributing to roughness perception. Here we extrapolated these ideas to our stationary single point stimulation paradigm, where participants did not freely explore a surface but instead received a single vibration waveform at the hand. Waveforms with richer high frequency content, such as a sawtooth, should be judged as rougher than smoother waveforms with limited high frequency energy, such as a sinusoid. Likewise, waveforms with more irregular or less predictable temporal structure should elicit higher roughness ratings than more regular periodic signals. Building on these considerations, Experiment 3 tested the hypothesis that perceived roughness differs systematically across waveform types in both auditory and vibrotactile modalities, with stimuli containing greater high frequency content and greater temporal irregularity being judged as rougher.

Finally, we examined whether roughness perception was systematically related to affective judgments of pleasantness. In audition, increased roughness is strongly associated with dissonance and reduced pleasantness [27]. This relationship also appears in broader multisensory accounts of roughness [30]. In touch, a similar relationship has been reported, with tactile roughness negatively correlated with pleasantness across a range of textured surfaces [31]. We therefore hypothesized that, in both auditory and tactile modalities, stimuli judged as rough would also be rated as less pleasant. The Experiment 4 tests this hypothesis by asking participants to provide pleasantness ratings for the same set of vibrotactile and auditory stimuli used in the roughness and frequency experiments.

In summary, we investigate what musical information can be conveyed through a tactile vibration alone using a single actuator. Specifically, we focus on how tactile stimuli can represent three key musical dimensions: pitch, roughness, and pleasantness. First, participants were asked to judge the perceived frequency of sinusoidal and complex vibrations to evaluate the tactile equivalent of pitch perception (Experiments 1 and 2). Second, they rated the roughness of the same complex stimuli to explore whether vibration can encode qualities related to consonance and dissonance (Experiment 3). Finally, they evaluated the pleasantness of the stimuli to examine how tactile signals influence emotional and aesthetic responses and if it is negatively correlated with roughness (Experiment 4). These experiments aimed to capture both low level perceptual attributes and higher level affective responses, providing a comprehensive understanding of the tactile representation of musical information.

Our findings in normal-hearing participants may inform the design of vibrotactile displays that support access to selected music-relevant perceptual cues.

## II. Methods

### A. Participants

A total of 25 participants took part in the study (18 female, mean age = 28.3 years, range = 20–42). Twelve participated in the auditory group and 20 in the tactile group, with an overlap of 7 participants who took part in both studies.

All participants reported normal hearing and normal tactile sensitivity, and none reported neurological disorders. All provided informed written consent before participation. The study was approved by the Ethics Committee of the University of Geneva and conducted in accordance with the Declaration of Helsinki.

### B. Apparatus

The experiment was conducted in a quiet room in the basement of the University of Geneva. Participants were comfortably seated in a reclining chair equipped with two armrests. The auditory stimuli were presented directly through headphones (Sennheiser HD280 pro) at a comfortable level and were normalized using a standard loudness model [32], [33]. Tactile stimuli were delivered by instructing participants to lightly grasp a snooker ball with their left hand, using a grip similar to shifting gears (see Figure 1). This approach ensured maximum skin contact without applying any pressure or force. To optimize the activation of mechanoreceptors, the entire hand was tested rather than just a single finger. The armrests were positioned to fully support the weight of the participants’ arms. The snooker ball was connected to a piezoelectric stack actuator (P-841K191, Physik Instrumente) that provided controlled stimulation. The displacement amplitude was calibrated using a laser vibrometer (Polytec OFV 503) and is expressed in decibels relative to 1 micrometer (dB re 1 µm). To eliminate any potential auditory cues from the actuator with the tactile stimuli, participants wore sound-attenuating headphones that played 84 dB SPL white noise.

**Fig. 1.**
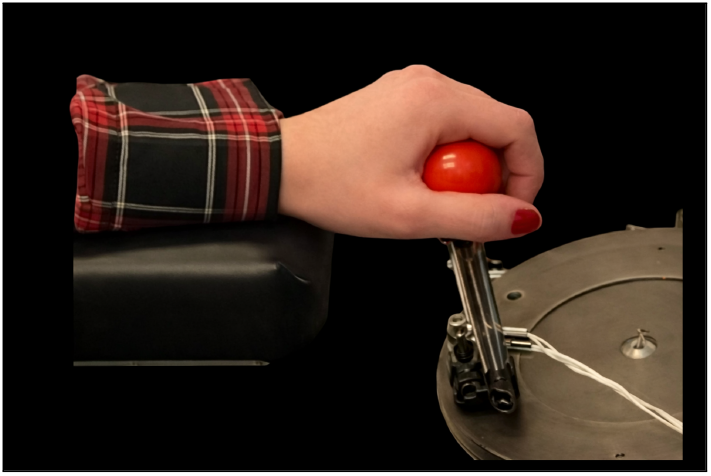
Vibrotactile stimulation setup. The participant’s forearm was rested on a padded support while the hand grasped a snooker ball coupled to a piezoelectric stack actuator (PZT).

### C. Stimuli

All tactile stimuli were 500 ms in duration, with 10 ms raised cosine onset and offset ramps to minimize transient artifacts. Experiment 1 used sinusoidal vibrations at 40, 80, 160, 200, 280, 400, 480, 600, and 750 Hz. Experiments 2 to 4 used six waveform types with three fundamentals, *F*_0_ = 52, 73, 98 Hz, corresponding to the musical notes *G#1, D2*, and *G2*. These *F*_0_ values span less than one octave, avoid simple harmonic relations (octave or fifth), and remain sufficiently low so that most energy of the harmonics fall within the vibrotactile perceptual range (below 1 kHz).

The six types of waveforms were selected to produce strongly different time domain waveforms and spectral envelopes, while preserving the same repetition rate, as confirmed by their autocorrelation functions (Figure 2). A sinusoid served as the baseline. A sawtooth, with a sharper temporal onset, tested whether transient like features enhance tactile pitch. To dissociate temporal sharpness from harmonic density, we included a harmonic complex with harmonics attenuated by 3 dB per octave and randomized phase, producing a smoother waveform while maintaining a rich harmonic structure. To assess whether tactile pitch can arise from periodicity in the absence of energy at the fundamental, analogous to the auditory missing fundamental phenomenon [25], [34], we created two filtered variants, Missing *F*_0_ (fundamental removed) and Missing *F*_0_*/F*_1_ (fundamental and first harmonic removed). Finally, an amplitude modulated, AM, stimulus (carrier 500 Hz, modulation rate = *F*_0_*s*) tested whether a clear low frequency temporal periodicity can drive tactile pitch even when most spectral energy is concentrated at higher frequencies, well above the fundamental. The carrier was set to 500 Hz so that, even for the highest modulation rate, the lowest prominent sideband (402 Hz) remained well above the effective bandwidth of low-frequency mechanoreceptors (Meissner corpuscles), while the upper sideband (598 Hz) remained clearly perceptible.

**Fig. 2.**
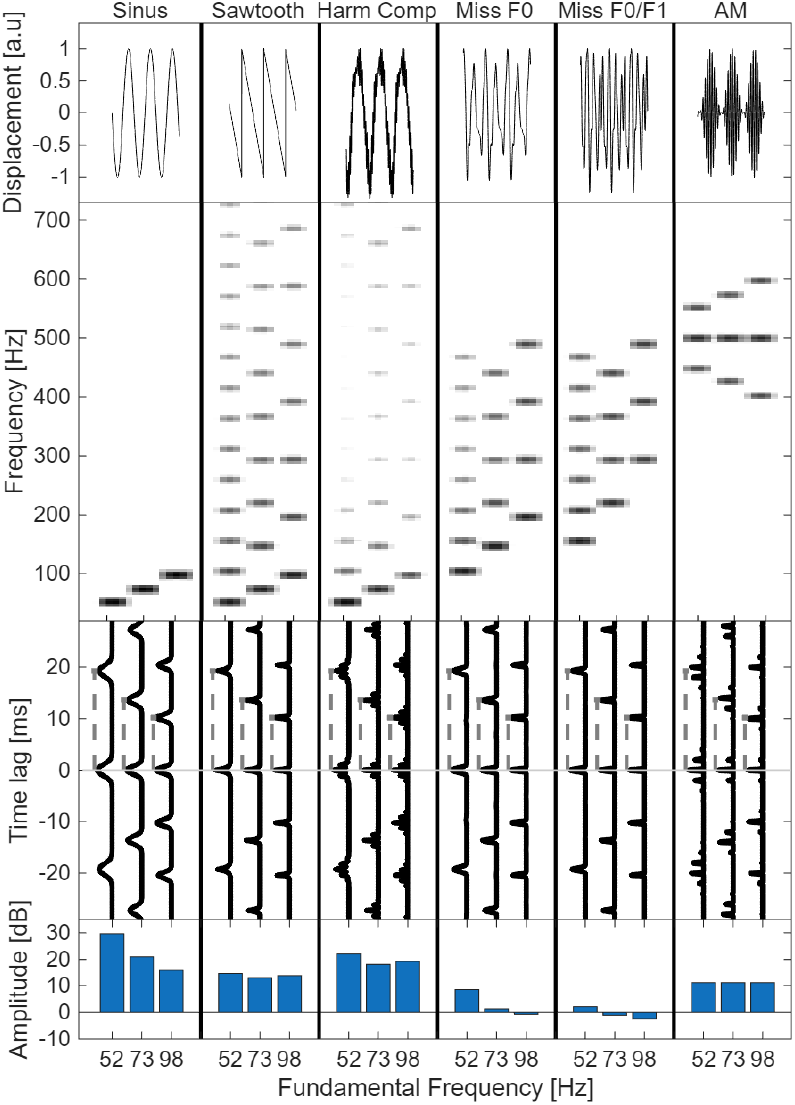
Physical representation and perceptual calibration of the 18 tactile stimuli grouped by waveform and fundamental frequency *F*_0_. The top row shows each waveform in the time domain. The second row presents spectrograms illustrating the frequency composition of each stimulus. The third row displays autocorrelation functions, demonstrating that all stimuli share the same periodicity despite differences in waveform and spectral envelope. The fourth row shows peak–to–peak displacement amplitudes in dB relative to 1 *µ*m, set by an expert panel to achieve equal perceived tactile intensity across stimuli.

Some of these waveforms, in particular the sawtooth and the missing fundamental variants with sharp edges and strong high order content, are difficult to reproduce accurately with a conventional voice coil actuator. Because a voice coil behaves as a spring mass system, its mechanical compliance and resonances tend to smooth rapid transitions and distort the intended time domain shape. We therefore used a piezoelectric actuator to achieve the required bandwidth and temporal fidelity. The resulting surface motion was verified with a laser displacement sensor, confirming that the measured output closely matched the target waveforms, see Figure 3 for representative recordings.

**Fig. 3.**
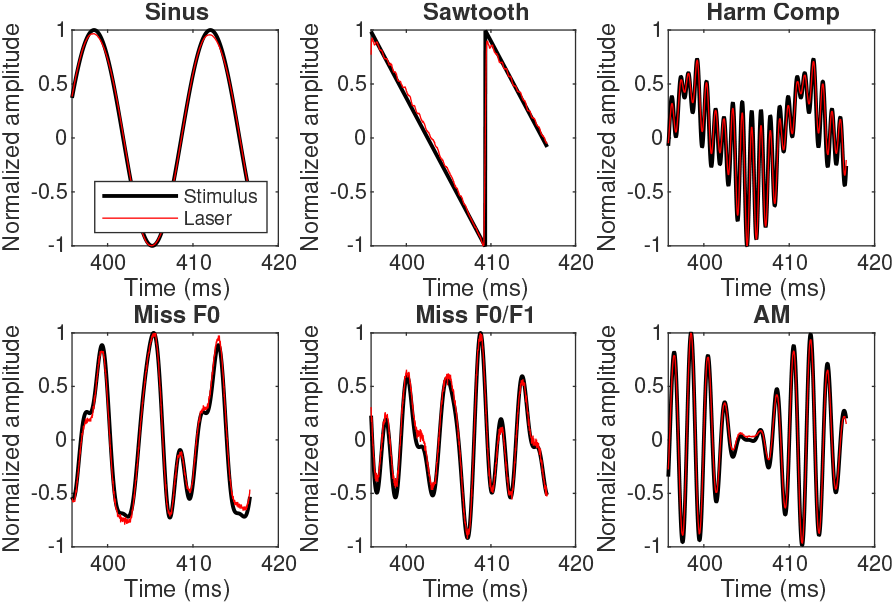
Laser vibrometry validation of the piezo actuator output. For each stimulus waveform, the commanded input (normalized) is shown together with the laser measured surface displacement (normalized), demonstrating that the piezo reproduces the intended temporal structure with high fidelity, including waveforms with sharp transitions and strong high order spectral content that are typically distorted by spring mass dynamics in voice coil actuators.

The amplitudes of the stimulus and harmonic components were calibrated in a three stage procedure. First, a group level equal perceived intensity (EPI) curve for sinusoidal vibrations was measured in a preliminary experiment with 19 participants who did not take part in the main study. Using a method of adjustment [35], participants matched the perceived intensity of sinusoidal test frequencies to a fixed sinusoidal reference vibration at 280 Hz played at *−*10 dB [re 1 µm]. This reference was chosen because 280 Hz lies within a highly sensitive frequency region, and it also corresponds to the mid range frequency tested in Experiment 1. Matched levels were expressed in dB relative to the reference and averaged across participants to yield an EPI curve specifying, for each frequency, the level required to evoke the same perceived intensity as the reference (see Figure 4).

**Fig. 4.**
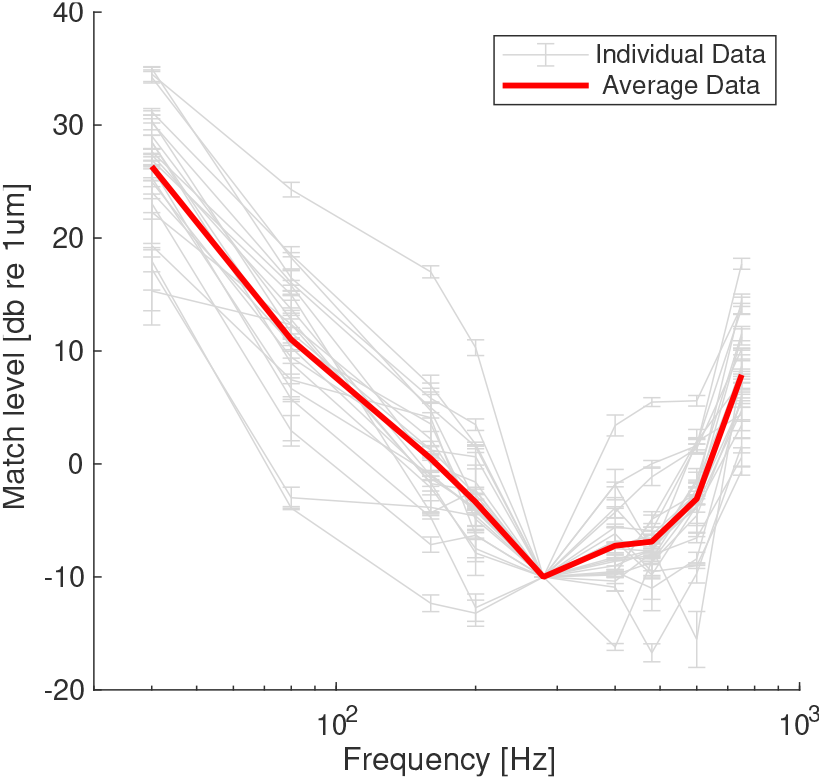
Equal perceived intensity (EPI) curves for tactile stimuli. Grey lines represent individual participants, while the thick red line represents the median response.

Second, for harmonic, missing *F*_0_, and related complex waveforms, the EPI curve was used as a spectral weighting filter during waveform synthesis. Each harmonic component was first assigned its nominal amplitude rule, for example 3 dB/octave for the harmonic complex, then additionally re-weighted according to the EPI value at that component frequency. This ensured that partials in highly sensitive regions contributed less physical amplitude and partials in less sensitive regions contributed more, thereby reducing the tendency for harmonics around a few hundred hertz to be disproportionately salient.

Third, after EPI-based spectral weighting, overall stimulus levels were set by an additional perceptual matching experiment conducted by an expert panel composed of 6 participants who did not participate in the main experiments. For each waveform type and each fundamental frequency, participants adjusted the global amplitude to match the perceived intensity of a sinusoidal reference at the same *F*_0_. The resulting waveform-specific global level offsets were averaged across participants and applied as fixed corrections in the main experiments (see Figure 2, last row).

Finally, to account for inter-individual differences in tactile sensitivity, sinusoidal stimulus levels in Experiment 1 were individualized using detection thresholds measured during the screening session. For each participant, the deviation of their threshold from the group typical threshold, evaluated at the calibration reference condition, was converted to dB and applied using a half gain rule, such that the participant specific correction was set to half of the threshold deviation. This reduced sensitivity driven differences while preserving the intended across frequency EPI profile.

### D. Tasks

First, a tactile threshold screening was conducted prior to the experiment to ensure that all participants had sensitivity within the normal range. Thresholds were measured using sinusoidal vibrations at frequencies between 40 and 750 Hz, delivered via the piezoelectric actuator. A two-interval forced-choice procedure with a 1-up 2-down adaptive rule was used to estimate the detection threshold for each frequency.

Then, all participants completed Experiments 1–4 within the modality group to which they were assigned, tactile only (n= 13), auditory only (n= 5), or both modalities (n= 7). Participants who completed both modalities always performed the auditory sessions second, at least two months after the tactile sessions, to minimize potential transfer or anchoring effects of explicit auditory pitch judgments on subsequent tactile frequency judgments.

Each experiment comprised a series of trials. On each trial, a single stimulus was presented, after which participants provided an absolute magnitude estimate. Participants were free to choose any numeric scale, with the instruction to remain internally consistent across the full session and across blocks. Responses were given verbally and entered by the experimenter.

Three tasks were assessed in separate blocks, perceived frequency (Experiments 1–2), roughness (Experiment 3), and pleasantness (Experiment 4). The perceived frequency task was designed to quantify vibrotactile pitch. We used the term “perceived frequency” instead of “pitch,” or its French equivalent “hauteur,” because in pilot testing participants were confused by these terms and could not readily relate them to a tactile sensation. Throughout the manuscript, we use “perceived frequency” and “pitch” interchangeably.

We explained this term as “how fast the vibration oscillates,” and illustrated it with a bouncing ball animation (1 to 8 bounces) to emphasize the perceived speed of repeated impulses. For roughness, we explained the adjective as the irregularity or harshness of the vibration, and illustrated it using physical sandpaper samples ranging from smooth to coarse, which participants were instructed to use only as a qualitative reference for “smooth versus rough,” not as a scale to be matched. For pleasantness, participants rated overall liking, supported by a graded set of facial expressions from unhappy to happy.

Each experiment was conducted as a single block lasting approximately 10 minutes. In Experiment 1, the 9 stimuli were each repeated 15 times and presented in randomized order, yielding 135 trials. In Experiments 2, 3, and 4, the 18 stimuli were each repeated 5 times, again in randomized order. Before each block, participants completed a short training run using the same task and stimulus set, terminated after 20 trials. Training trials were excluded from analysis and served only to familiarize participants with the task and help them establish an internal numeric scale.

The order of blocks was randomized across participants to counterbalance potential learning, fatigue, and order effects. A single session of approximately 2 hours was scheduled to complete all stages of the protocol, including instructions, threshold, training runs, breaks, and the experimental blocks.

For analysis, repeated trials for each stimulus and participant were first aggregated using the geometric mean. Because absolute magnitude estimation allows idiosyncratic numeric scales across participants, responses were then normalized by an anchor value within each participant. In Experiment 1, we fit the participant’s frequency judgment function with a 4th order polynomial and extracted the predicted value at 200 Hz, all judgments were divided by this value. In Experiments 2–4, judgments were normalized by dividing by the geometric mean response to the 72 Hz sinusoidal anchor stimulus.

## III. Results

### A. Experiment 1: Vibrotactile pitch induced by a sinusoidal stimulus

In Experiment 1, participants were asked to rate the perceived frequency of sinusoidal vibrations in the auditory and tactile modalities.For auditory stimuli, perceived frequency judgments increased linearly with the logarithm of stimulus frequency for all participants, showing minimal interindividual variability (Figure 5, top panel). To quantify how perceived frequency scales with physical frequency, we followed the classic approach used to derive the Mel scale [24] and modeled judgments as a linear function of log frequency. A linear mixed effects model fit to these judgments, with log frequency as a continuous fixed effect and participant as a random intercept, yielded a slope of 0.83 (*df* = 1, *p* < 0.001, *R*^2^ = 79%), reflecting strong sensitivity to frequency changes.

**Fig. 5.**
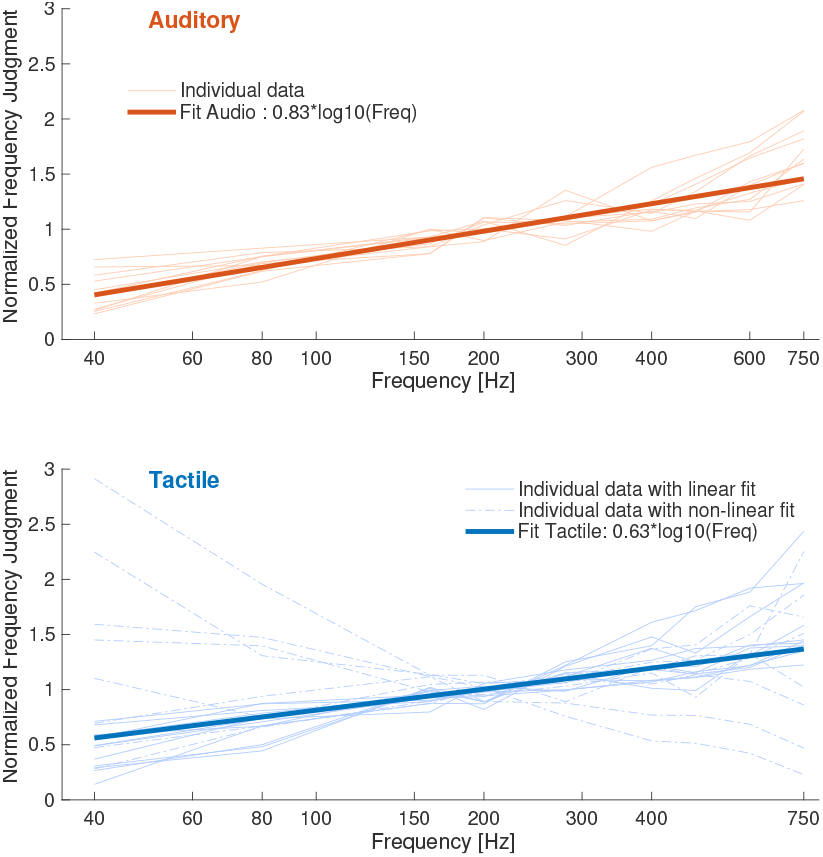
Perceived frequency judgments for sinusoidal stimuli across all participants. The upper panel displays individual responses for auditory stimuli (thin lines). The lower panel presents individual responses for tactile stimuli, where participants best fit by a positive linear regression are shown with thin continuous lines, while those best fit by other models are represented with thin dashed lines. The overall linear fit for both stimuli are depicted with thick lines.

Tactile stimuli also yielded a significant linear relationship, albeit with a shallower slope of 0.63 (*df* = 1, *p* < 0.001, *R*^2^ = 20%). When pooling data across modalities and including presentation mode as a categorical factor, the mixed linear model revealed significant main effects of presentation mode (*F*(1, 284) = 17.9), log of frequency (*F*(1, 284) = 133.49), as well as a significant interaction between the two (*F*(1, 284) = 17.5), all with *p* < 0.001. This interaction indicates a significant reduced sensitivity to frequency variations in the tactile condition compared to audition (Figure 5, bottom panel).

To explore the inter-individual differences, we fitted both linear and quadratic models to each participant’s responses and conducted ANOVAs to determine the most appropriate fit. The significance level *α* was adjusted for multiple comparisons using 0.05 divided by the number of participants per condition. All auditory responses best fit a positive linear model, with slopes ranging from 0.43 to 1.23. Tactile responses were varied: 13 out of 20 participants exhibited a positive linear fit (continuous lines, Figure 5), four favored a quadratic fit, two showed no clear relationship (null model), and one participant demonstrated a negative linear relationship, (dashed lines, Figure 5). The underlying reasons for these different behaviors in the tactile condition remain unclear. We examined variables such as gender, musical expertise, age, and familiarity with tactile tasks, but none provided a significant explanation.

When the linear mixed effects model was re-estimated while restricting the analysis to participants exhibiting a positive linear fit in the tactile condition, the ANOVA revealed a significant main effect of log frequency (*F*(1, 212) = 378.14, *p* < 0.001), but no effect of presentation mode (*F*(1, 212) = 0.18, *p* = 0.67) and no presentation mode × log frequency interaction (*F*(1, 212) = 0.15, *p* = 0.70). In this subset, the estimated slope remained shallower (0.72) but did not differ significantly between tactile and auditory conditions.

Taken together, these results partially support our first hypothesis, showing that when stimulus intensity is controlled, perceived frequency increases with physical frequency in both modalities, but only among participants who exhibited a positive linear fit in the tactile condition.

### B. Experiment 2: Vibrotactile pitch induced by a complex stimulus

In Experiment 2, participants rated the perceived frequency of complex periodic stimuli differing in waveform and fundamental frequency (illustrated in Figures 2 and 3). Despite individual variations, the median judgments increased with frequency for both auditory and tactile stimuli, except for the AM auditory stimuli (see Figure 6). Waveform type notably influenced auditory pitch perception, consistently resulting in higher pitch judgments for complex stimuli compared to sinusoidal stimuli of identical periodicity. This effect, known as the octave error, likely arises from greater energy in higher harmonics, influencing the perceived tone height [25].

**Fig. 6.**
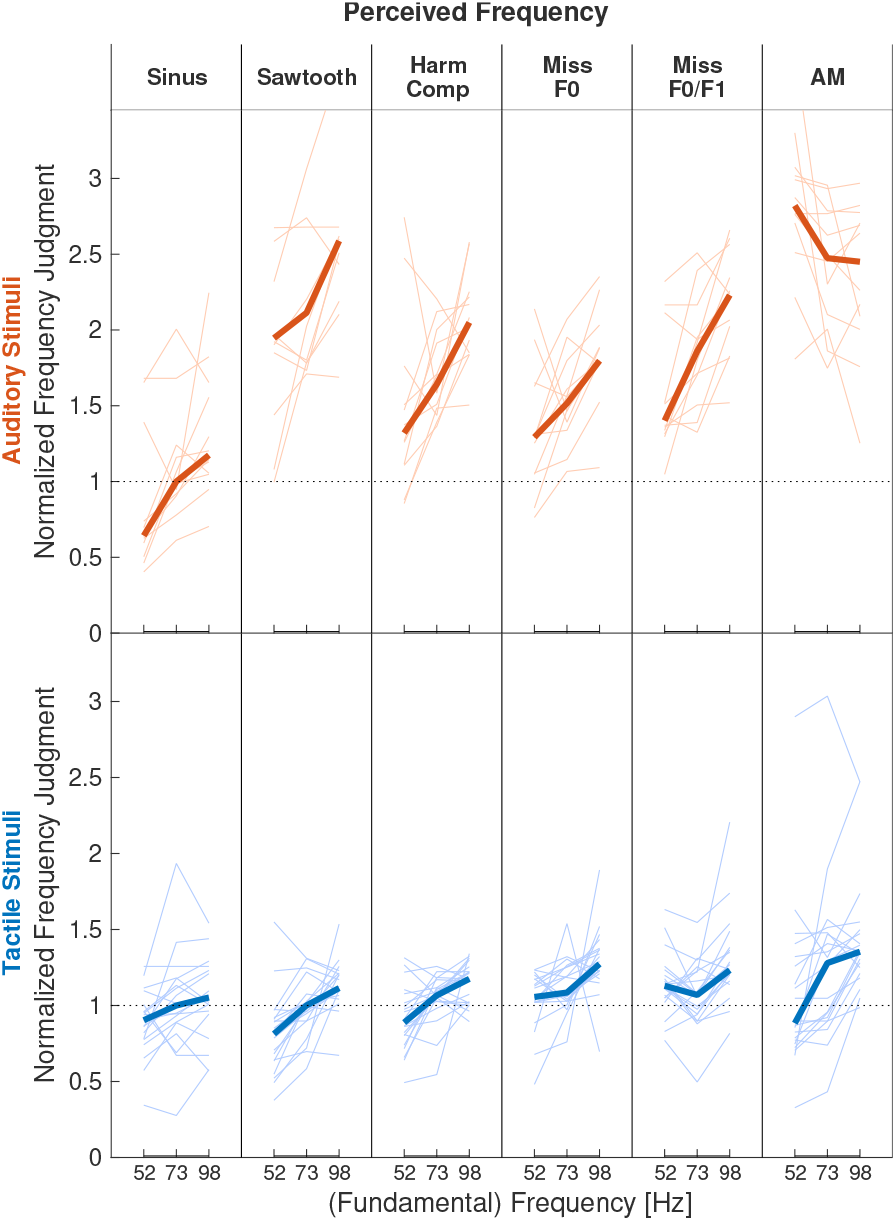
Results of Experiment 2. Normalized perceived frequency judgments for complex auditory (top panel) and tactile (bottom panel) stimuli as a function of *F*_0_. Thin lines indicate individual participant responses, thick lines represent the median judgments across participants. Perceived frequency increased monotonically with *F*_0_ in both auditory and tactile conditions. However, waveform-dependent differences were pronounced in the auditory modality, while tactile judgments showed little sensitivity to waveform type.

A two way repeated measures ANOVA on *auditory perceived frequency judgments* showed significant main effects of Waveform (*F*(5, 55) = 9.82, *p* < 0.0001, *R*^2^ = 37%), and Frequency (*F*(2, 22) = 9.93, *p* = 0.0008, *R*^2^ = 3%) with a significant interaction (*F*(10, 110) = 6.60, *p* < 0.0001, *R*^2^ = 3.2%). Post hoc Bonferroni analyses revealed that sinusoidal stimuli were judged significantly lower than the harmonic complex (*p* = 0.0003), Missing *F*_0_ (*p* = 0.0088), Missing *F*_0_*/F*_1_ (*p* = 0.0012), and AM (*p* = 0.0002). Additionally, AM stimuli were judged significantly higher than harmonic complex (*p* = 0.0212), Missing *F*_0_ (*p* = 0.0007), and Missing *F*_0_*/F*_1_ (*p* = 0.0035). Missing *F*_0_ and Missing *F*_0_*/F*_1_ also differed significantly (*p* = 0.0127).

A similar two way repeated measures ANOVA on *tactile perceived frequency judgments* showed significant main effects of Waveform (*F*(5, 95) = 4.27, *p* = 0.0014, *R*^2^ = 11%), and Frequency (*F*(2, 38) = 38.37, *p* < 0.0001, *R*^2^ = 10%), with a significant interaction (*F*(10, 190) = 2.17, *p* = 0.02, *R*^2^ = 2%). Post hoc Bonferroni analyses showed that sinusoidal stimuli were judged significantly lower than Missing *F*_0_ (*p* = 0.0355). Sawtooth stimuli were judged significantly lower than Missing *F*_0_ (*p* = 0.0032), and Missing *F*_0_*/F*_1_ (*p* = 0.0029). Additionally, harmonic complex stimuli were judged significantly lower than Missing *F*_0_ (*p* = 0.0256). Notably, the sinusoidal stimuli were not significantly lower than Missing *F*_0_*/F*_1_, in accordance with the effect of the missing fundamental in tactile perception.

Comparing the auditory and tactile ANOVAs via effect sizes, *R*^2^, we observed that auditory pitch perception was primarily driven by Waveform explaining 37% of variance, compared to only 3% for frequency. In contrast, tactile pitch perception showed similar modest influences from Waveform and Frequency each accounting for about 10% of variance.

While we hypothesized that waveform shape and spectral composition would have a strong effect on vibrotactile pitch even when periodicity was held constant, the effect size pattern shows a clear modality difference. In audition, pitch judgments were dominated by Waveform, whereas in touch Waveform and Frequency contributed comparably and modestly. This suggests that vibrotactile pitch, at least under our single point stimulation conditions, relies primarily on temporal periodicity cues, with a more limited contribution of waveform dependent, timbre like information than in the auditory domain. In other words, the hypothesis is only partially supported, waveform does matter for tactile pitch, but it does not exert the strong, dominant influence observed for auditory pitch.

Given that waveform only modestly affected the *magnitude* of vibrotactile pitch, we introduced the complementary hypothesis that waveform may instead modulate *pitch salience*. If vibrotactile pitch is partly mediated by Pacinian (PC) afferents, which are particularly sensitive to rapid changes in skin indentation and function as effective change detectors, then a waveform with an abrupt pressure transition, such as a sawtooth, should yield a more salient pitch percept than a smooth waveform such as a sinusoid. We therefore hypothesized that waveforms with a sharp rise should lead to a more salient perception of vibrotactile pitch. To test this, we compared the effect of Frequency on the sinusoidal and sawtooth stimuli using a planned contrast analysis. Frequency indeed had a significantly stronger effect, i.e., a steeper slope, on the sawtooth stimulus compared to the sinusoidal stimulus (*p* = 0.0239), supporting this salience hypothesis.

### C. Perception of Roughness

In Experiment 3, participants rated the perceived roughness of the complex stimuli in the auditory and tactile modalities. Roughness judgments varied systematically with waveform type in both modalities, while fundamental frequency had a limited effect (see Figure 7, left panel).

**Fig. 7.**
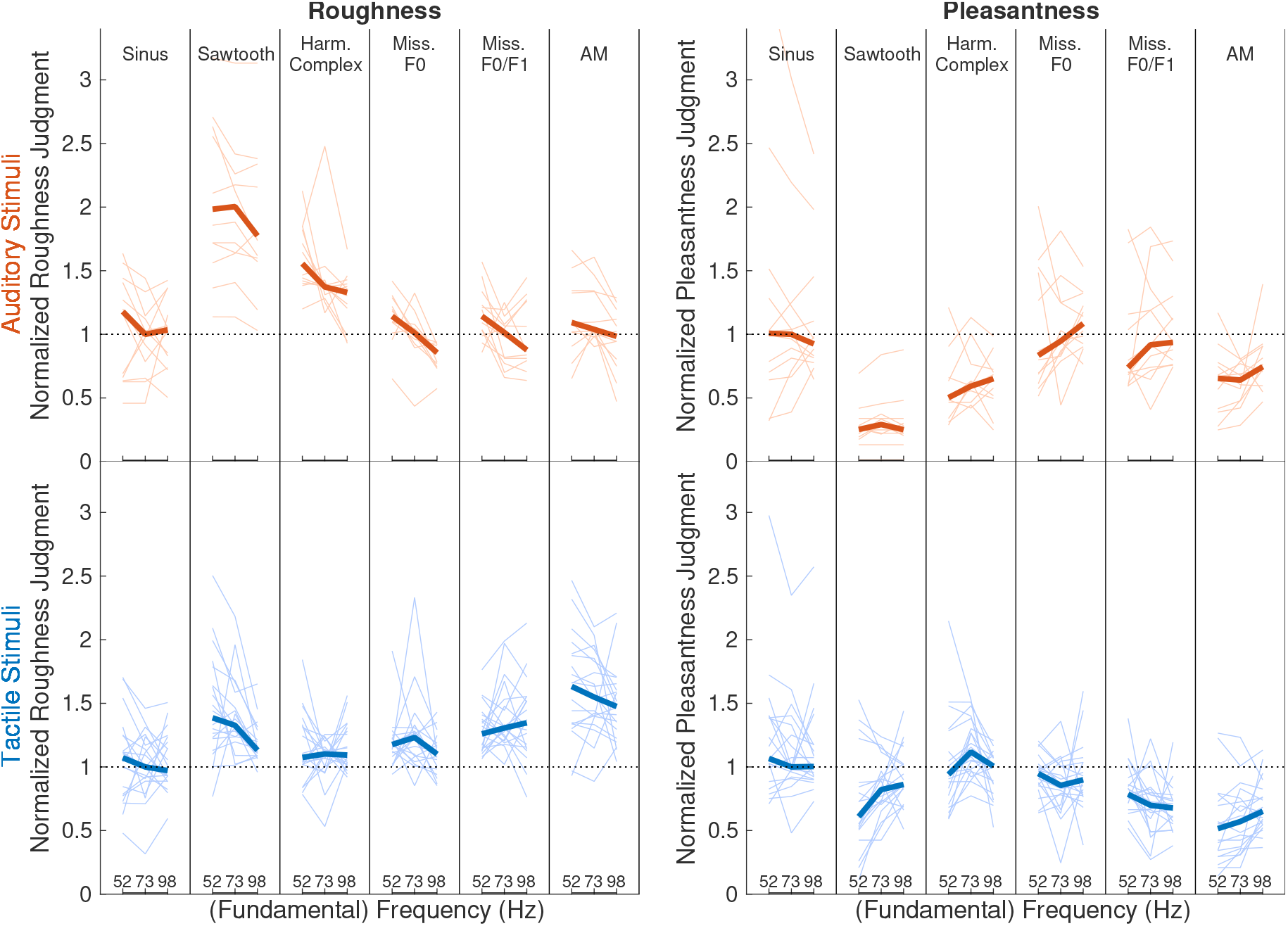
Left panel: roughness ratings. Right panel: pleasantness ratings. Normalized roughness and pleasantness ratings are shown for six waveform types: sinusoidal, sawtooth, harmonic complex, Missing *F*_0_, Missing *F*_0_*/F*_1_, and AM, at three base frequencies (52, 73, and 98 Hz), for auditory stimuli (upper panels, orange) and tactile stimuli (lower panels, blue). Thin lines indicate individual participant responses, thick lines represent the median judgments across

Consistent waveform effects emerged across both modalities. For the auditory stimuli, a two way repeated measures ANOVA on *roughness ratings* revealed a significant main effect of Waveform (*F*(5, 55) = 18.49, *p* < 0.0001, *R*^2^ = 52%), and Frequency (*F*(2, 22) = 43.11, *p* < 0.0001, *R*^2^ = 2%), but no significant interaction (*F*(10, 110) = 1.59, *p* = 0.1174). Post hoc pairwise comparisons, Bonferroni corrected, showed that the sawtooth stimulus was rated significantly rougher than the sinusoid (*p* = 0.0200), Missing *F*_0_ (*p* = 0.0034), Missing *F*_0_*/F*_1_ (*p* = 0.0059), and AM (*p* = 0.0070), as predicted by the model. In addition, the harmonic complex was rated rougher than the sinusoid (*p* = 0.0426), AM (*p* = 0.0097), and both Missing *F*_0_ conditions (*p* = 0.0004 and *p* = 0.0017). As expected, *F*_0_ had only a minor influence on roughness perception.

For the tactile stimuli, a significant main effect of Waveform was found on *roughness ratings*(*F*(5, 95) = 22.14, *p* < 0.0001, *R*^2^ = 28%), along with a weak and significant interaction with Frequency (*F*(10, 190) = 2.95, *p* = 0.0018, *R*^2^ = 3%), but no main effect of Frequency alone (*F*(2, 38) = 1.77, *p* = 0.1835). Post hoc comparisons revealed that AM stimuli were perceived as significantly rougher than all other waveforms (all *p* < 0.01 excluding the sawtooth *p* = 0.0466). The sawtooth waveform was rated rougher than the sinusoid and harmonic complex, but smoother than the AM. The harmonic complex was judged smoother than both the Missing *F*_0_*/F*_1_ /*p* = 0.0055), and sawtooth (*p* = 0.0008), and the sinusoidal waveform was consistently rated as the smoothest, significantly smoother than Missing *F*_0_*/F*_1_ (*p* = 0.0067), and sawtooth (*p* < 0.0001).

The AM waveform was perceived as significantly rougher than all stimuli. This emphasizes the critical role of waveform irregularities in the perception of tactile roughness. In addition, the sawtooth was rated as the second roughest. Rather than frequency content alone, tactile roughness appears particularly sensitive to disruptions of regularity and sharp temporal fluctuations. This aligns with our proposed hypothesis that temporal waveform irregularity and abrupt amplitude transients, rather than harmonic density per se, dominate tactile roughness perception under single point stationary stimulation [28].

We conclude that tactile roughness can be systematically controlled by modifying the vibration waveform, with minimal influence from frequency. If we aim to use vibration to convey musical emotion, we must examine whether tactile roughness contributes to feelings of unpleasantness, as has been observed in the auditory domain [30]. In the next experiment, we tested whether tactile roughness is negatively correlated with tactile pleasantness.

### D. Experiment 4: Perception of Pleasantness

In Experiment 4, participants rated the perceived pleasantness of the same set of stimuli. Pleasantness ratings differed across waveform types and were negatively correlated with roughness in both auditory and tactile conditions (see Figure 7, right panel). Pleasantness ratings for the complex stimuli revealed a significant effect of Waveform, especially for auditory stimuli. For the auditory stimuli, a two way repeated measures ANOVA on *pleasantness ratings* indicated a significant effect of Waveform (*F*(5, 55) = 9.54, *p* < 0.0001, *R*^2^ = 37%), while there was no main effect of Frequency (*F*(2, 22) = 0.66, *p* = 0.5276), nor interaction effect between Waveform and Frequency (*F*(10, 110) = 0.93, *p* = 0.5093). Post hoc analyses further reveal that participants rated the sawtooth waveform as significantly less pleasant than all the other waveforms, sinusoidal (*p* = 0.041), harmonic complex (*p* = 0.0013), Missing *F*_0_ (*p* = 0.0011), Missing *F*_0_*/F*_1_ (*p* = 0.003), and AM (*p* = 0.0074). The harmonic complex was judged as less pleasant than the Missing *F*_0_ (*p* = 0.0016). Interestingly, within the other waveforms, AM was judged as less pleasant than both Missing *F*_0_ (*p* = 0.0186 and *p* = 0.033).

For tactile stimuli, the ANOVA also shows a significant influence of Waveform on pleasantness (*F*(5, 95) = 15.15, *p* < 0.0001, *R*^2^ = 29%), and a significant, though weaker, interaction effect with Frequency (*F*(10, 190) = 3.93, *p* < 0.0001, *R*^2^ = 4%), while Frequency alone does not significantly impact pleasantness ratings, (*F*(2, 38) = 0.02, *p* = 0.9779). Notably, the AM tactile stimulus received significantly lower pleasantness ratings compared to sinusoidal (*p* = 0.0017), sawtooth (*p* = 0.0327), harmonic complex (*p* < 0.0001), and Missing *F*_0_ (*p* = 0.0004) stimuli. The sawtooth waveform was judged as less pleasant than the harmonic complex (*p* = 0.0247). As for the audio stimuli, the two tactile stimuli that were judged as more rough, were also judged less pleasant. Additionally, Missing *F*_0_*/F*_1_ was rated as less pleasant than the sinusoidal (*p* = 0.0117), harmonic complex (*p* = 0.0002), and Missing *F*_0_ (*p* = 0.0009).

Because judgments of pleasantness are highly individual, we examined the relationship between roughness and pleasantness at the level of each participant. For each person, we computed the correlation between their roughness and pleasantness ratings across the full set of stimuli. Correlations were calculated over the 18 stimuli, yielding 17 degrees of freedom.

For the auditory stimuli, a significant negative correlation (*p* < 0.05), was observed in 10 out of 12 participants (Figure 8), and for the tactile stimuli, in 12 out of 20 participants (Figure 9). All significant correlations were negative, except for P46 with the tactile stimuli, indicating that rougher sounds or vibrations are generally judged as less pleasant.

**Fig. 8.**
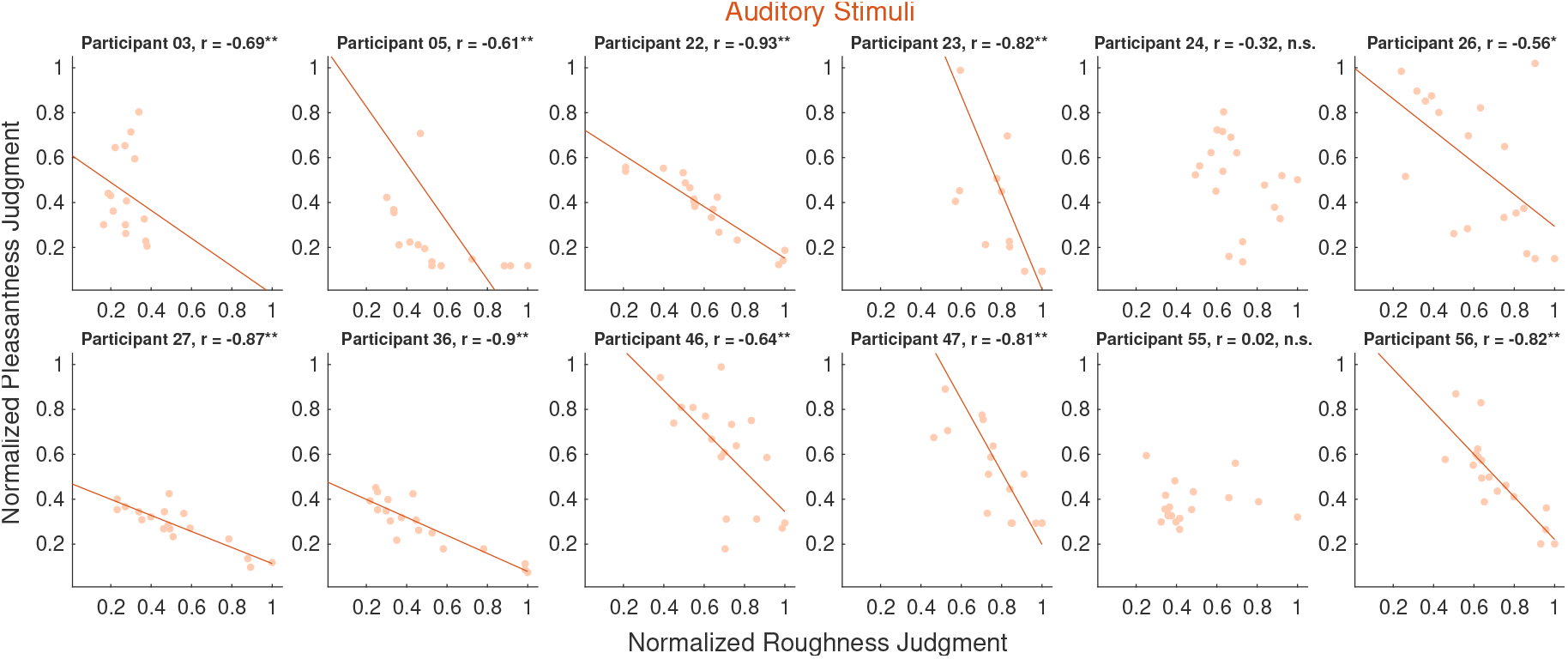
Correlation between roughness and pleasantness ratings for auditory stimuli. Each panel shows the data of an individual participant. Dots represent single stimulus ratings, and straight lines indicate significant regression fits.

**Fig. 9.**
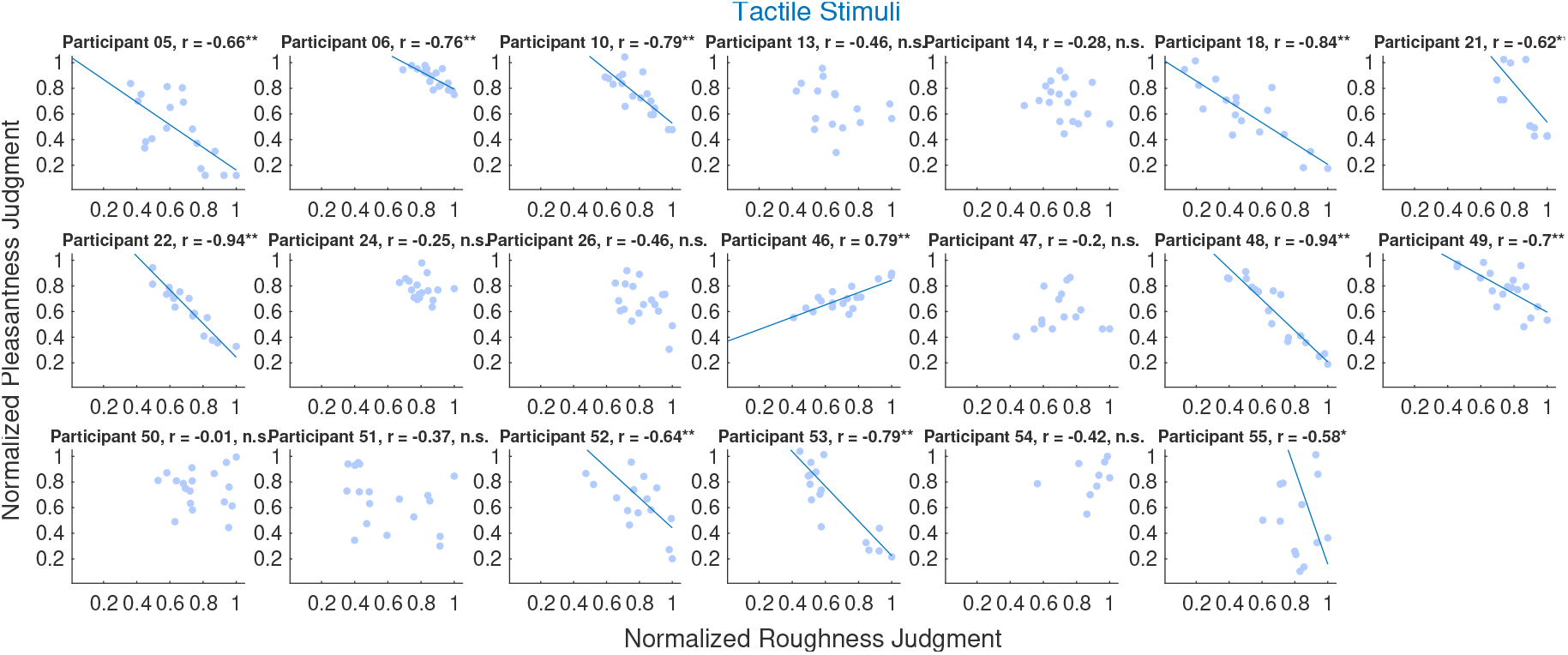
Correlation between roughness and pleasantness ratings for tactile stimuli. Each panel shows the data of an individual participant. Dots represent single stimulus ratings, and straight lines indicate significant regression fits.

This experiment confirms that, as in audition, tactile roughness can influence affective judgments. Stimuli that were perceived as rougher were also rated as less pleasant, and this relationship was robust at both the group and individual level.

## IV. Discussion

We explored how core musical features, pitch, roughness, and pleasantness, can be conveyed through tactile vibration. Our findings highlight a central difference between auditory and tactile processing: the capacity for spectral decomposition.

In audition, the basilar membrane mechanically separates incoming sound into narrow frequency bands, enabling precise encoding of both spectral and temporal features. In contrast, vibrotactile signals are processed largely through two specialized frequency channels, the Meissner, 5 to 50 Hz, and Pacinian corpuscles, 40 to 1000 Hz, which offer limited spectral resolution [36]. As a result, tactile perception relies more heavily on the fast temporal envelope than on detailed spectral content. Nevertheless, our data indicate that vibrotactile stimuli can convey some musical information and these limitations are crucial to consider when designing an effective haptic device for conveying music.

### A. The perception of pitch through vibrotactile stimulation

We observed a monotonic relationship between the physical frequency of vibrotactile stimuli and their perception, Figure. 5. This suggests that changes in melody contour can indeed be encoded through variations in the frequency of tactile stimuli. However, the slope describing the relationship between the stimulus frequency and its perception was smaller than in the auditory domain. Practically, this means that larger frequency differences are required in the tactile domain to achieve perceptual changes equivalent to those in the auditory domain. For example, while a 6% frequency difference in auditory stimuli is sufficient to produce a perception of one semitone, the distance between two adjacent notes on a piano, an 8% difference is necessary for tactile stimuli. Consequently, direct translation of audio signals to tactile stimuli would result in a compressed perception of melody contour. This finding aligns with previous studies that have demonstrated significantly larger just noticeable differences for tactile frequencies compared to auditory frequencies [12], [19], [20].

### B. The effect of waveform on the perception of vibrotactile pitch

We hypothesized that a sinusoidal vibration might not be the optimal stimulus to convey pitch. Therefore, our study explored how waveform differences affect vibrotactile pitch perception. Specifically, we found that stimuli with abrupt discontinuity, such as sawtooth waves, were more sensitive to frequency changes than simpler and smoother waveforms like sinusoids. Although our Experiment on complex perceived frequency was limited to a relatively narrow frequency range, the data suggest that the slope relating physical frequency to perceived frequency might be steeper for the sawtooth waveforms than for the sinusoidal. We propose that the enhanced sensitivity observed results from the efficient detection by Pacinian corpuscles of more salient changes in the stimulus.

To illustrate this hypothesis, we modeled the responses of tactile neurons using TouchSim, a somatosensory neuron activation simulation [37]. The simulations demonstrated that sinusoidal waveforms induced neuronal firing at various phases, accompanied by minor temporal jitter, Figure 10. In contrast, the sawtooth waveform elicited strong neural activation at two specific moments corresponding precisely to the waveform’s abrupt changes, Figure 10. It is important to note a fundamental difference in how auditory and tactile systems respond to oscillatory stimuli. In the auditory system, inner hair cells are activated only during positive pressure displacement of the basilar membrane, while negative displacement does not trigger activation. Since sound waves are oscillatory, every positive pressure phase is followed by a negative one; thus, auditory hair cells typically activate once per cycle, at frequencies below the phase locking limit [38]. In contrast, Pacinian corpuscles respond to both positive and negative pressure displacements, resulting in two activations per cycle [39]. This difference in activation mechanisms is clearly observed in our model (Figure 10).

**Fig. 10.**
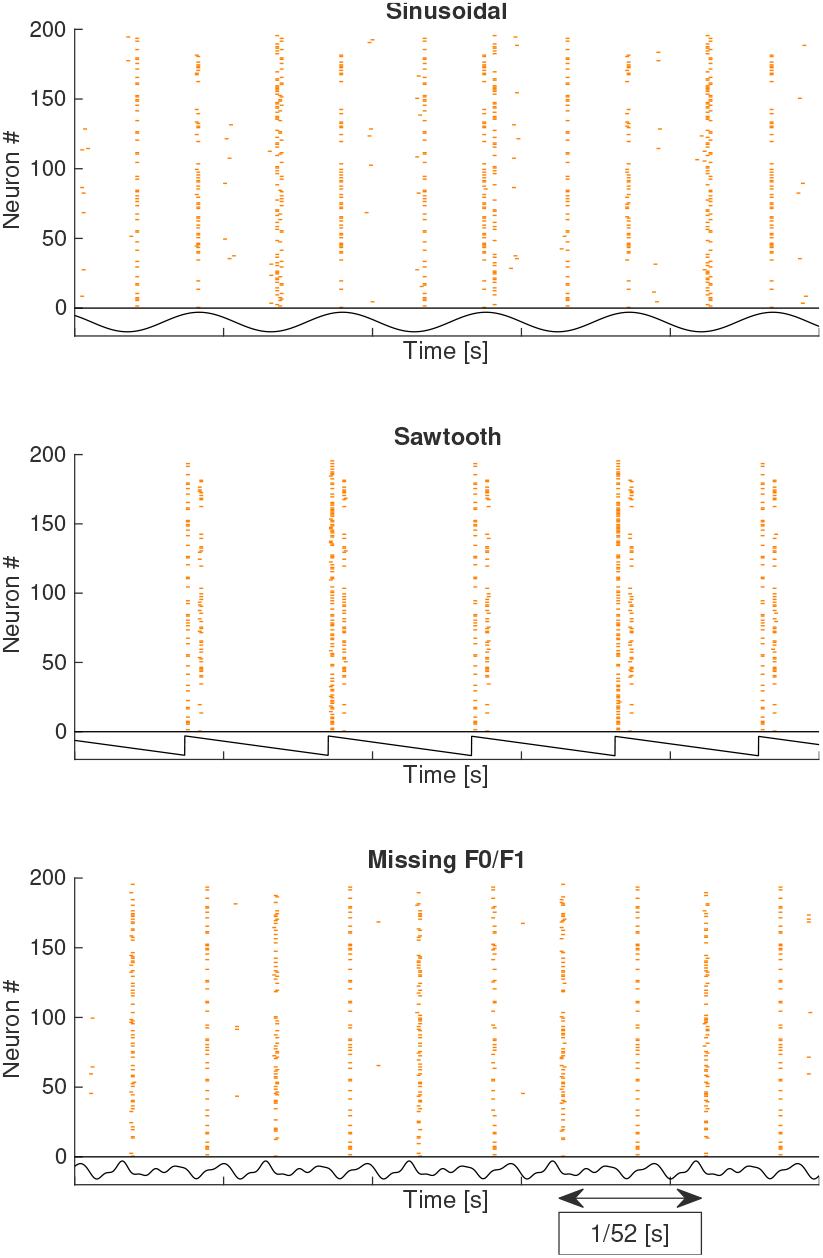
Modeled neuronal responses to a) sinusoidal, b) Sawtooth, c) Missing F0/F1 waveforms with a (fundamental) frequency of 52 Hz, using TouchSim [37]. The figure shows the activation pattern of 200 PCs along the different waveforms. Notably, the Sawtooth waveform elicited pronounced neural activations corresponding to abrupt slope transitions, while waveforms lacking fundamental frequency energy still produced activation patterns reflecting their periodicity.

It is also important to note that since all stimuli were matched in perceived intensity, they differed significantly in maximal displacement. For instance, at 52 Hz, the sinusoidal waveform required a displacement amplitude 5.6 times greater than the sawtooth waveform to achieve the same perceptual intensity. Therefore, to convey pitch effectively using a simple waveform, the sawtooth not only produces a more salient percept but also requires substantially smaller actuator displacement. However, generating sawtooth mechanical movements is technically more challenging compared to sinusoidal waveforms and cannot be achieved with simple actuators like ERMs.

### C. The effect of the missing fundamental is perceivable in the tactile domain

Our data provide support for the missing fundamental phenomenon in the tactile domain. Specifically, waveforms lacking energy at the fundamental frequency, Missing *F*_0_ and Missing *F*_0_*/F*_1_, were perceived with comparable or nearly comparable frequencies to sinusoidal vibrations. Despite their distinct spectral envelopes, all tested stimuli shared identical periodicity, as confirmed by autocorrelation analysis, see Figure 2. Similarly, the modeled neuronal responses, Figure 10, demonstrated repetition periods in synchrony to sinusoidal stimuli. Analogous to findings in the auditory domain, these results suggest that tactile pitch perception primarily depends on stimulus periodicity rather than the physical presence of the fundamental frequency itself [25], [26]. To our knowledge, these results are consistent with a tactile correlate of the auditory missing fundamental, supporting an envelope periodicity account.

This finding contributes to the growing evidence of perceptual parallels between auditory and vibrotactile systems. Furthermore, leveraging the missing fundamental effect offers a promising strategy to create lower-frequency vibrotactile pitches without the need to generate large-displacement, low-frequency vibrations, which can be difficult and energy-intensive to produce based on the vibrotactile equal-loudness contour. By employing this technique, achievable also through amplitude-modulated waveforms [40] lower tactile pitches could be produced efficiently with significantly reduced amplitude requirements.

### D. The physiological mechanism of vibrotactile pitch

One proposed mechanism for vibrotactile pitch perception is the ratio hypothesis [41], which suggests that pitch is determined by the relative activation of Meissner and Pacinian corpuscles. These mechanoreceptors have distinct frequency tuning: in humans, Meissner corpuscles are most sensitive to vibrations between 5 and 50 Hz, while Pacinian corpuscles are tuned to the higher range from 40 to 800 Hz, with peak sensitivity at 250 Hz [36]. According to the ratio hypothesis, lower frequency stimuli would primarily activate Meissner corpuscles and higher frequency stimuli would increasingly recruit Pacinian corpuscles. Vibrotactile pitch, under this framework, would be encoded by the relative contribution of each receptor population.

If this hypothesis held, we would expect a systematic increase in perceived frequency across stimuli that shift energy toward higher frequencies, for example, from sinusoidal to Missing *F*_0_, then Missing *F*_0_*/F*_1_, and finally to AM stimuli, which contain only high frequency components within the Pacinian range. Furthermore, as soon as the frequency content of the stimuli lies entirely above the Meissner range, such that only Pacinian corpuscles are stimulated, any further change in spectral content should not affect perceived frequency. That is, pitch perception should plateau once Meissner activity is minimal or absent.

However, our results do not align with this prediction. We observed similar pitch percepts for sinusoidal and Missing *F*_0_*/F*_1_ stimuli, despite large differences in their spectral content and presumed receptor activation patterns. More critically, AM stimuli, composed entirely of high frequency carriers within the Pacinian range, showed clear changes in perceived frequency as a function of modulation frequency. This contradicts the expectation that pitch should remain constant once only the Pacinian channel is engaged. A similar pattern was observed in Experiment 1, where changes in frequency continued to affect its perception even in a range that would exclusively stimulate Pacinian corpuscles.

While these findings argue against a strict implementation of the ratio hypothesis, we cannot fully rule it out. The hypothesis was originally formulated based on responses to sinusoidal stimuli, and it is possible that, with more complex signals, the skin and somatosensory system act as a low pass filter, effectively extracting the temporal envelope. This would shift the relevant energy toward the fundamental frequency or modulation rate, potentially explaining the observed alignment in pitch percepts across stimuli with very different spectra. Nonetheless, the failure of perceived frequency to plateau once Meissner input is presumably absent challenges the idea that vibrotactile pitch is determined solely by the relative activation of these two afferent populations.

While these findings argue against a strict implementation of the ratio hypothesis, we cannot fully rule it out. The hypothesis was originally formulated based on responses to sinusoidal stimuli, and it is possible that, with more complex signals, the skin and early somatosensory processing act as a low pass filter or envelope detector, effectively extracting the temporal envelope periodicity. Under this simpler account, our “missing *F*_0_” conditions could be interpreted as *envelope-based periodicity perception* rather than a direct tactile analogue of the auditory missing fundamental, because higher harmonics may not be fully available at the skin. This would shift the relevant energy toward the fundamental period or modulation rate, potentially explaining the observed alignment in perceived frequency across stimuli with very different spectra. Nonetheless, the failure of perceived frequency to plateau once Meissner input is presumably absent challenges the idea that vibrotactile pitch is determined solely by the relative activation of these two afferent populations.

### E. Perception of tactile roughness and pleasantness

Roughness, an important attribute linked to musical dissonance, exhibited significant effects of waveform but was highly idiosyncratic across participants. Unlike auditory roughness, which relies on temporal fluctuations, tactile roughness is often influenced by spatial texture cues, such as fine surface details experienced during hand movement [30]. In our experiment, however, roughness was conveyed purely through temporal cues, as participants interacted with a smooth snooker ball as the contact surface. This setup created a novel and unfamiliar task, likely contributing to the response variability.

Nevertheless, the inter participant variability could be attributed to consistent personal preferences, as roughness judgments showed a negative correlation with pleasantness, mirroring findings in auditory perception where rougher stimuli are generally rated as less pleasant [30], [31]. Waveforms influenced both roughness and pleasantness rating similarly. Waveforms with sharper edges, like the sawtooth and irregularity like the AM stimuli, were perceived as both rougher and less pleasant compared to smoother waveforms like the sinusoidal or harmonic complex, Figure 7. High frequency components, often linked to harshness in auditory perception, appeared to play a similar role in tactile perception [30].

### F. The perception of the fast envelope instead of the fine structure

In auditory science, signals can be conceptually decomposed into two complementary components: the temporal envelope and the temporal fine structure. The temporal envelope refers to the slowly varying amplitude modulation of the waveform, capturing broad fluctuations that guide perceptual attributes such as rhythm, speech intelligibility, and overall loudness. Conversely, the temporal fine structure encompasses the rapid, cycle by cycle fluctuations within the waveform, crucial for precise pitch perception, source localization, and discerning complex auditory scenes. Cochlear implant users, who rely predominantly on envelope information due to technological limitations in encoding fine structure, often exhibit significant deficits in perceiving pitch nuances and understanding speech in noisy environments [26].

Similarly, our tactile stimuli can be analyzed using this decomposition. However, the distinction between envelope and fine structure in tactile signals depends on the chosen temporal resolution. In this study, we adopted a high temporal resolution, defining a fast temporal envelope that accurately follows rapid amplitude transitions, such as those in a sawtooth waveform. Our results consistently demonstrate the critical role of this fast temporal envelope in tactile perception. Specifically, the prominence of pitch and roughness perception observed with sharp edged waveforms like the sawtooth highlights the envelope’s significance. Moreover, the tactile equivalent of the auditory missing fundamental phenomenon observed in our experiments further supports the conclusion that vibrotactile pitch perception primarily relies on the periodicity conveyed by the fast envelope rather than on detailed fine structure elements.

### G. Limitations of the study

This study provides insights into how tactile stimulation can convey selected music relevant perceptual cues, but several limitations must be acknowledged.

First, the equal intensity setting was critical to control for the interaction between frequency and intensity in the tactile domain [23], [42]. Individual calibration ensured that stimuli were perceived at comparable intensity levels across participants. However, this approach is difficult to translate directly to real world musical contexts, where intensity and frequency vary dynamically over time. It therefore remains unclear how well such static calibrations would generalize to continuous music, or whether adaptive, context dependent adjustments would be required to maintain perceptual consistency.

Second, all participants had normal hearing, therefore we cannot infer how these tactile judgments relate to music perception in people with hearing loss, including cochlear implant users. In addition, stimulation was delivered through a single contact configuration, participants grasped a single interface with the hand, and responses may differ for other body sites, coupling conditions, or contact forces. Moreover, all tactile stimuli were delivered using a single actuator at a single point. While this design isolates the effect of waveform structure on basic perceptual attributes, it does not capture the richer spatiotemporal cues available in multi actuator haptic displays.

Third, the stimuli were presented in isolation, which does not reflect the complexity of music. In audition, multiple tones can be perceived concurrently, supported by stream segregation mechanisms that build on the cochlea’s decomposition of sound into frequency selective channels. Tactile perception lacks an equivalent spectral decomposition, and it is therefore uncertain whether a single point vibration source could convey multiple tactile pitches simultaneously. This limitation highlights the likely need for multi actuator approaches, or alternative encoding strategies, to approximate polyphonic musical structure.

Fourth, the complex waveform set was intentionally restricted to low fundamental frequencies (52, 73, 98 Hz), therefore conclusions about waveform dependent changes in perceived frequency, roughness, and pleasantness should be limited to this low *F*_0_ range. Although we did not find a main effect of frequency on roughness and pleasantness, previous studies have reported relationships between perceived roughness and the peak frequency content of texture induced vibrations during sandpaper scanning [28]. In addition, Barbosa Escobar and Wang, using a single wrist band actuator, linked vibration frequency to crossmodal taste associations, with lower frequencies (around 50 Hz) associated with more pleasant taste descriptors and higher frequencies (around 100 Hz) with more sour descriptors [43]. Relatedly, although Experiment 1 included an auditory frequency matching task spanning 40–750 Hz, Experiments 2–4 focused on low *F*_0_ stimuli by design, so the auditory–tactile comparisons primarily inform this region.

Finally, judging the perceived frequency of a vibration is an unusual task. Unlike auditory frequency, which has clear everyday relevance through speech and music, tactile frequency judgments are rarely made explicitly, which likely contributed to the substantial inter individual variability observed here. More ecological tasks, for example discrimination within musical sequences or mapping to categorical rhythmic or timbral descriptors, will be needed to link these controlled psychophysical judgments more directly to real world musical experience.

### H. Impact for developing a tactile device

For the tactile stimuli, we found that many participants showed similar response slopes to those in the auditory condition, suggesting that the perception of musical pitch can extend to touch. This supports the idea that tactile melodies are feasible and that vibration based devices could be used for music perception. Although the tested frequency range, 40 to 750 Hz, is narrower than the full auditory range of 27.5 Hz to 4186 Hz in Western music, a meta analysis of 10,000 piano solo pieces revealed that most notes played fall within 262 to 523.5 Hz, *C*4 to *C*5, which is within the tactile sensory spectrum [44].

To perceive the higher notes, outside the tactile range, music could be pitch shifted down by one or two octaves [13], [45]. However, this approach assumes that participants will perceive the differences between notes with the same magnitude as in the auditory modality. When considering the average slope across all participants, it was significantly reduced. Additionally, just noticeable differences for frequency are larger than the auditory just noticeable difference, even among experienced musicians [12]. This implies that a tactile melody may not be perceived in the same way as an auditory melody, but rather as a compressed version. To mitigate this effect, one potential solution is to stretch the melody temporally, as tested by Aker et al. [11]. The degree of stretching would need to be carefully calibrated so that stimuli presented an octave apart are perceived as the same musical note.

Our findings provide valuable insights for designing tactile devices aimed at conveying musical information. First, the use of complex waveforms, such as harmonic complexes or sawtooth waveforms, can enhance the clarity of perceived frequency by leveraging the tactile system’s ability to interpret periodicity and harmonic structure. Additionally, since roughness and pleasantness were influenced by waveform and frequency, devices could dynamically adjust these parameters to evoke specific emotional or musical effects, such as creating tension through rougher waveforms, such as AM stimuli, or resolving it with smoother, more pleasant stimuli, such as harmonic complex.

The observed negative correlation between roughness and pleasantness highlights the importance of balancing these factors when designing tactile feedback. For example, using sinusoidal or harmonic complex waveforms could prioritize pleasantness for general music playback, while sharper waveforms, such as the sawtooth, might be reserved for encoding dissonance or tension. Furthermore, frequency range adjustments, such as pitch shifting music down by one or two octaves, could align the signal with the most salient frequencies in the tactile modality, while preserving the overall structure of musical content.

Finally, future tactile devices should account for individual differences in tactile perception by incorporating customizable calibration routines. Such routines could optimize stimulus intensity, waveform selection, and frequency presentation based on user specific thresholds and preferences. Incorporating multi actuator systems could also help overcome the limitations of single point stimulation by enabling more complex spatial patterns, approximating the richness of auditory stream segregation in music.

## V. Conclusions

We explored the potential of tactile stimuli to convey key aspects of musical information, including pitch, roughness, and pleasantness. By investigating the tactile perception of sinusoidal and complex waveforms, we assessed whether tactile devices can replicate critical elements of music.

Our findings suggest that tactile stimuli can effectively convey certain musical dimensions. Approximately two thirds of participants perceived frequency in a manner similar to auditory pitch, with a linear relationship between perceived frequency and the logarithm of the presented frequency. Additionally, waveforms with harmonic content were found to enhance the clarity of frequency perception, highlighting the importance of waveform design in tactile pitch representation.

For roughness and pleasantness, individual variability was more pronounced, particularly in the tactile modality. While roughness judgments were influenced by frequency and waveform, these effects were idiosyncratic. However, the observed negative correlation between roughness and pleasantness aligns with auditory findings, suggesting that tactile roughness could serve as a proxy for dissonance in musical applications.

Despite these promising results, several challenges remain. The narrower frequency range of tactile perception, the reduced sensitivity to frequency changes, and the limited ability to present overlapping signals with single point actuators highlight the need for further advancements in tactile technology.

Finally, our findings reinforce the importance of distinguishing between the fast temporal envelope and the temporal fine structure in vibrotactile perception. Therefore, future tactile musical interfaces should focus on optimizing the fast envelope to effectively convey musical dimensions, thereby enhancing the tactile experience of music for hearing impaired users.

Taken together, these results in normal hearing participants provide quantitative constraints on how waveform structure shapes perceived frequency, roughness, and pleasantness in touch. This may support the development of vibrotactile strategies for transmitting selected music relevant cues, but translation to real world musical experience, and validation in hearing impaired groups, including cochlear implant users, are essential next steps.

## Acknowledgements

We would like to thank Alastair Loutit, Pablo Mavro cordatos, and Gregorio Avila for their valuable comments and assistance with the study design. We are also grateful to the staff of the Brain and Behavior Laboratory at the University of Geneva for their support with the experimental setup and environment.

